# Inference of interactions between chromatin modifiers and histone modifications: from ChIP-Seq data to chromatin-signaling

**DOI:** 10.1101/010132

**Authors:** Juliane Perner, Julia Lasserre, Sarah Kinkley, Martin Vingron, Ho-Ryun Chung

## Abstract

Chromatin modifiers and histone modifications are components of a chromatin-signaling network involved in transcription and its regulation. The interactions between chromatin modifiers and histone modifications are often unknown, are based on the analysis of few genes, or are studied *in vitro*. Here, we apply computational methods to recover interactions between chromatin modifiers and histone modifications from genome-wide ChIP-Seq data. These interactions provide a high-confidence backbone of the chromatin-signaling network. Many recovered interactions have literature support; others provide hypotheses about yet unknown interactions. We experimentally verified two of these predicted interactions, leading to a link between H4K20me1 and members of the Polycomb Repressive Complexes 1 and 2. Our results suggest that our computationally derived interactions are likely to lead to novel biological insights required to establish the connectivity of the chromatin-signaling network involved in transcription and its regulation.

## Introduction

Transcription and its regulation are facilitated by a complex interplay between various molecular players such as transcription factors, chromatin modifiers (CMs), histone modifications (HMs) and RNA polymerase II (Pol II). Together these components form a chromatin-signaling network (1) whose signaling activity affects the transcriptional and the chromatin state of a particular genomic region. Thus, it is not surprising that the presence of certain HMs at the promoter or the gene body coincides with the transcriptional status of the corresponding gene (2, 3). This close link is further substantiated by the finding that there is even a quantitative relationship between HM levels and the steady state level of mRNAs (4-6).

HMs are closely linked to the transcriptional process but their functional role in transcription remains largely unknown. On one hand HMs may modulate the stability of nucleosomes or the chromatin conformation (7) and thereby directly interfere with Pol II recruitment or processivity. On the other hand, HMs may play an indirect role by recruiting CMs to well-defined regions of the genome. Thus, because histones are firmly bound to DNA, HMs may restrict the signaling activity to certain genomic features such as enhancers and promoters.

The activity of the chromatin-signaling network leads to co-localization of HMs and CMs on the genome, which can be determined by Chromatin Immunoprecipitation followed by sequencing (ChIP-Seq; (8-10)). Accordingly, clustering HM and CM ChIP-Seq data identifies patterns of co-localized HMs and CMs, which can be associated with genomic features like enhancers and promoters (11). The co-localization pattern specific to e.g. promoters unravels those CMs and HMs that constitute the building blocks of the underlying chromatin-signaling network. However, such an analysis is unlikely to identify the specific interactions between CMs and HMs.

Recently, two approaches, one based on Bayesian Network inference (12) and the other on a maximum entropy framework (13), have been proposed to infer chromatin-signaling networks in *Drosophila melanogaster*. Both approaches require discrete data. This, however, involves difficult decisions on optimal decision thresholds. To circumvent these problems we use the ChIP-Seq levels directly and infer a human chromatin-signaling network. We construct this network drawing on two complementary philosophies. We model each HM level as a weighted linear combination of the CM levels and select those CMs that have the most consistent quantitative information about the HM level using Elastic Nets (14). This approach accounts for interactions induced by correlations between CMs, but is not able to remove interactions induced by correlations between HMs. Consequently, we prune the so-derived candidate chromatin-signaling network by computing sparse partial correlation networks (15), which is aimed to identify direct interactions between HMs and CMs accounting for correlations between CMs and HMs.

## Results

### HMs and CMs hold redundant information about gene expression

As both, HMs and CMs, are components of the chromatin-signaling network involved in transcription and its regulation, both should contain information about gene expression. To test this idea we used linear regression models to predict gene expression values from HM or CM levels at promoters in the human K562 cell line. The HM levels explain 76% of the variance in gene expression (Figure 1A), which is similar to the results from earlier work (4-6). The CMs capture 75% of the variance in gene expression (Figure 1B). The good predictive performance confirms that both HMs and CMs contain extensive information about gene expression.

**Figure 1.**
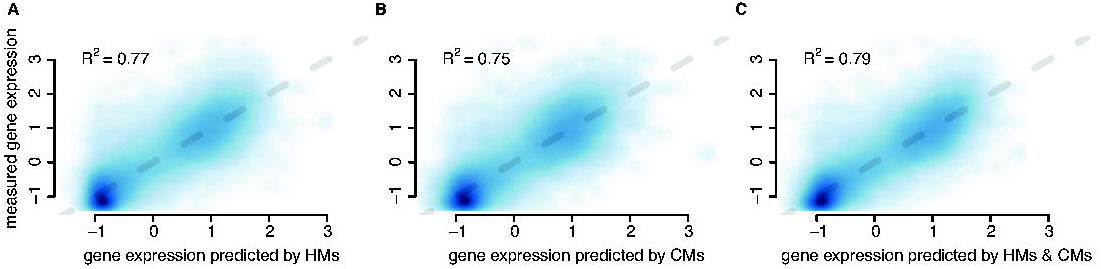
HMs and CMs hold redundant information about gene expression. Scatterplots with the predicted gene expression by HMs (A), CMs (B) and both (C) on the x-axis and the measured gene expression (CAGE-tags) on the y-axis. The blue color indicates the densities of points, the darker the denser. The grey dashed line indicates identity. In the left upper corner of each plot the coefficient of determination (R2), i.e. the variance in the gene expression measure explained by the model, is indicated.

If HMs and CMs reflect the same chromatin-signaling network, both should contain redundant information about gene expression such that combining them should yield only a marginal increase in the predictive power. Indeed, using both, CMs and HMs, improves the explained variance in gene expression only by 3% (4%) compared to using only HMs (CMs) at the expense of a higher model complexity (Figure 1C). Thus, these findings support that CMs and HMs jointly constitute a chromatin-signaling network involved in transcription and its regulation.

### CM levels predict HM levels and vice versa

Given that CMs and HMs are coupled together by the chromatin-signaling network, the levels of CMs should contain information about the HM levels and *vice versa*. To test this idea we separated the HMs from the CMs and modeled each group of variables using the other. For each HM we built simple linear regression models using 10-fold cross-validation (CV) and predicted the HM level based on a weighted combination of the CM levels. For all HMs the models account for at least 50% of the variance in the HM or CM level (Figure 2A). For H3K9ac, H3K4me3, H3K4me2 and H3K27ac the model explains even more than 85% of the variance, which is close to the agreement between biological replicates (Supplementary Figure S1). The high explanatory power of CMs for these four HMs suggests that many CMs interact with these HMs. Indeed, roughly half of the CMs are known to interact with modifications of the H3K4, the H3K9, or the H3K27 residue (Supplementary Table S1).

**Figure 2.**
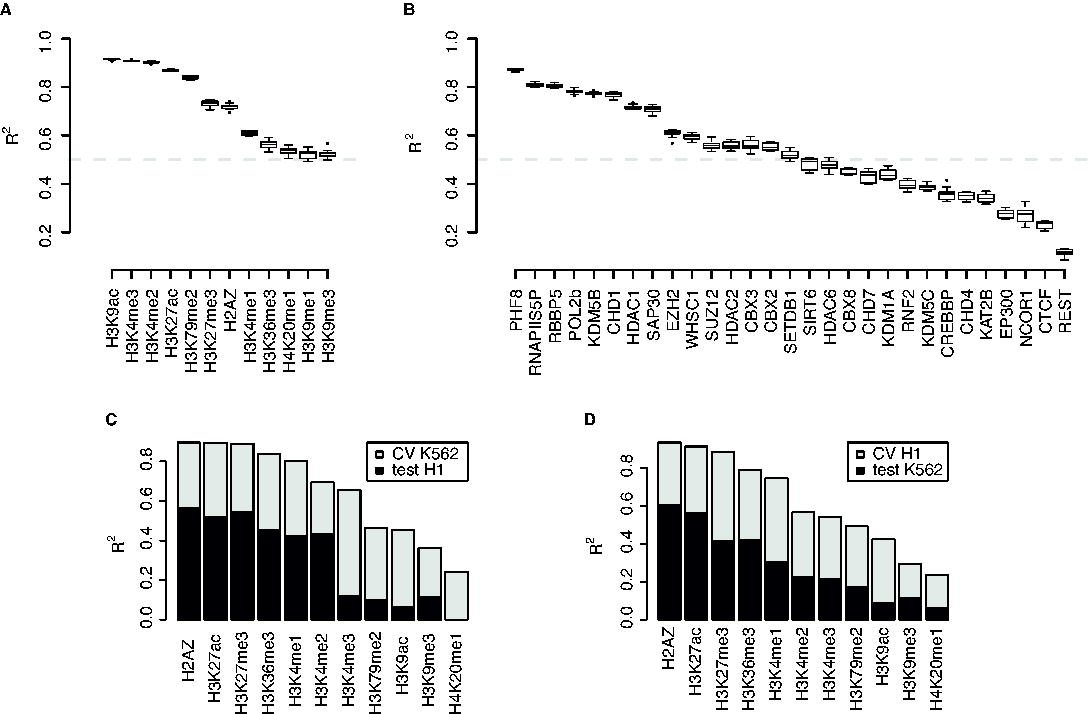
CM levels predict HM levels and vice versa. (A, B) Boxplots showing the range of coefficients of determination (R2) obtained by 10-fold cross-validation using CMs to predict HMs (A) and HMs to predict CMs (B). The boxes indicate the range of R2 values between the first and third quartile, the horizontal thick line indicates the median and the whiskers extend the range to 1.5 fold the range from the median to the lower and upper hinge of the box. R2 values outside this range are depicted as points. The dashed grey line indicates an R2 of 0.5. (C, D) Barplots showing the coefficients of determination (R2) obtained by training the model with data from the K562 cell line and testing it in the H1 cell line (C) and by training in H1 and testing in K562 (D). The total height of the bar indicates the average R2 obtained by 10-fold cross-validation in the training cell line, while the darker part indicates the R2 obtained by testing in the other.

We repeated this analysis by predicting CM levels from a linear combination of HM levels. For about half of the CMs the models account for over 50% of the variance (Figure 2B). Thus, for those well-predicted CMs the HMs in the data set cover the bulk of the recruitment mechanisms and enzymatic targets.

Under the assumption that the chromatin-signaling network is a common mechanism underlying transcription and its regulation, we expect that the contribution of a CM to the prediction of an HM in one cell type is similar in another cell type. Thus, given the regression model trained on the data from K562 cells we should be able to predict the HM levels in another cell type. We tested this using ChIP-Seq data for 14 CMs and 11 HMs in human embryonic stem cells (hESCs) that were also measured in the K562 cells. Indeed, the regression models learned from the data available for both cell types show good agreement (Figure 2C and D). The lower performance of the models when tested on the data from a different cell type is expected due to biological variation, e.g. different expression levels of the CMs. Thus, the quantitative effects of the interactions within the chromatin-signaling network are preserved suggesting a cell-type independent chromatin-signaling network involved in transcription and its regulation.

### From co-localization to interactions

We have shown that CM levels accurately predict HM levels and *vice versa*. We argued that the prediction accuracy depends on the expression and biochemical activities of the available CMs towards the HMs. To identify CM-HM pairs that are likely to interact with each other, we selected those CMs that contributed most to the prediction of an HM level. The most straightforward approach is to select those CM-HM pairs that show the highest pair-wise correlation. This has been done in recent work by clustering HMs and CMs into correlated subgroups based on their co-occupancy patterns (11).

There are groups of HMs and CMs that exhibit very high pairwise correlation (Supplementary Figure S2), suggesting that they are functionally related. However, within these groups no internal structure is visible, rendering an identification of interactions between the group members difficult. As CMs and HMs constitute a chromatin-signaling network, this high correlation is expected due to direct interactions between its components. However, high correlations could also be induced by other factors connecting the respective CM and HM. In general, the identity of these additional factors is not known, but we can account for those factors that are present in the dataset. Thus, we want to recover interactions between CMs and HMs that cannot be “explained away” by other variables in the dataset.

We recovered these interactions by applying a two-step procedure (see Methods). First, we used a regularized regression technique called “Elastic Net”, where the CMs are used to predict HMs, to select only CMs that are informative for the prediction of a HM. Moreover, in case of groups of strongly correlated CMs the members of these groups tend to remain all in the model or are removed together (14). This approach accounts for possible interactions induced by correlations within the CMs but does not take into account correlations between the HMs. This indicates that highly correlated HMs might be predicted by similar sets of CMs, while only certain CMs actually interact with specific HMs. Second, to remedy this situation we used a technique called “Sparse partial correlation networks” (SPCN; (15)), where the pairwise rank correlation between a CM and a HM is conditioned on all other variables in the data set. This method takes into account the correlation structure of both, CMs and HMs, and is conservative in proposing interactions. As a consequence, in groups of strongly correlated CMs and/or HMs, interactions may be explained away by individual members of the group (15). Thus, in the SPCN framework an identified interaction is likely to represent a direct interaction in the sense that it cannot be explained by other variables in the dataset. However, the failure to recover an interaction does not imply the absence of a biologically meaningful interaction. Within the SPCN framework some interactions between CMs and HMs arise from logical dependencies induced by sharing a common target. Thus, to recover interactions, we establish first the necessary condition that a CM is consistently highly predictive for an HM level by the Elastic Net approach and in a second step we prune those interactions that may be induced by correlations between the HMs using the SPCN approach. Thus, we focus only on interactions that are recovered by both methods. These interactions may originate from a direct function of the CM in setting, erasing or binding the HM but also from indirect interactions via unobserved CMs.

### Distinct sets of CMs associate with each HM

In the Elastic Net network each HM is linked to a different set of CMs indicating the different specificities of the CMs towards the individual HMs (Supplementary Figure S3A). The densest part of the network connects several CMs to the HMs H3K4me3, H3K9ac, H3K27ac and H3K79me2. The effect of the SPCN framework becomes most apparent on this dense cluster (compare Supplementary Figure S3A and B) where most of the interactions are resolved. It is important to note that the lack of a predicted interaction by the SPCN is not sufficient evidence to prove the absence of a biological relevant interaction. However, an interaction recovered by both approaches is likely to represent a true interaction between the CM and the HM.

### The chromatin-signaling network recovers biologically meaningful interactions

Many of the interactions identified by both Elastic Net and SPCN (Figure 3) are supported by published experimental evidence (Supplementary Note S1 and Supplementary Table S2), strengthening our confidence in the recovered interactions. For example, H3K27me3 has a positive interaction with members of the Polycomb Repressive Complex (PRC) 1 (CBX2 and CBX8; (16, 17)) and members of the PRC2 (EZH2 and SUZ12 (17)), as well as a negative interaction with Pol II phosphorylated at serine 5 (RNAPIIS5P).

**Figure 3.**
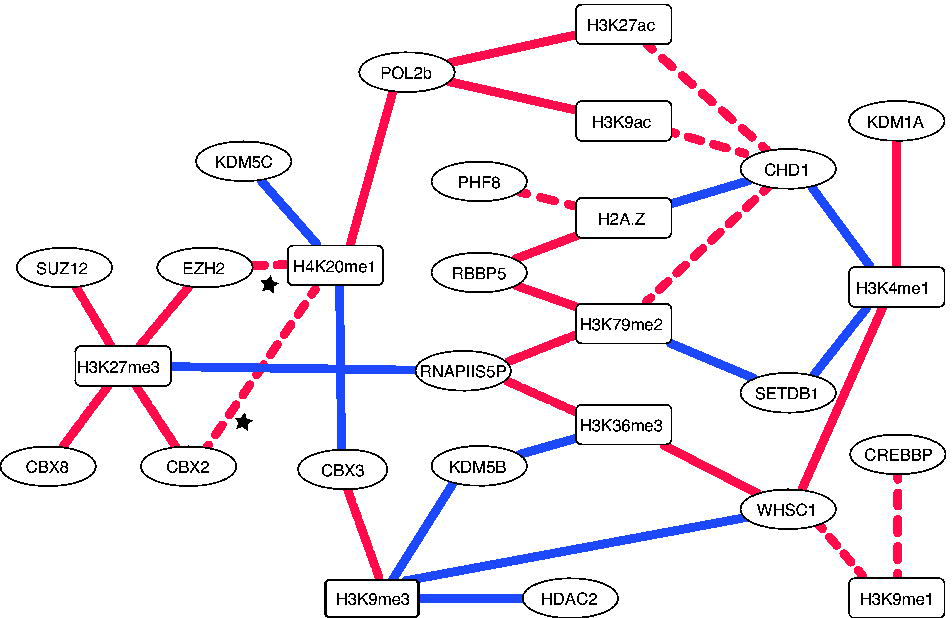
Chromatin-signalling network. Graphical representation of the interactions between CMs (circles) and HMs (squares). Shown are the interactions recovered both by the Elastic-Net and SPCN approach. Red lines indicate positive and blue lines negative interactions. The continuous lines indicate interactions with supporting evidence in the literature, while the dashed lines indicate interactions without supporting evidence. The stars indicate the two interactions confirmed in this study.

The interaction between H3K27me3 and EZH2 is direct, because EZH2 sets H3K27me3 (18-21). The interaction between H3K27me3 and SUZ12 may be direct, because it cannot be “explained away” by EZH2. However, EED which forms a trimeric complex together with SUZ12 and EZH2 binds H3K27me3 directly (22), and most likely explains the interaction between H3K27me3 and SUZ12 (23). The interaction between the PRC1 components CBX2 and CBX8 and H3K27me3 is direct, because CBX2 and CBX8 bind to H3K27me3 (20).

A negative interaction connects RNAPIIS5P and H3K27me3 in our network. The serine 5 phosphorylation of Pol II is mediated by the pre-initiation complex factor TFIIH (24-27) and is present in the initiating and the elongating form of Pol II (28). A role of H3K27me3 is to repress transcription, which is accompanied by low levels of initiating and/or elongating Pol II marked by serine 5 phosphorylation, explaining the negative interaction with RNAPIIS5P in our network.

Using H3K27me3 as an example, these results show that our approach identifies biological meaningful interactions between the members of PRCs and H3K27me3. If we did not have any prior information about the interactions between H3K27me3 and PRC, we would conclude that members of the PRCs are involved in setting and/or reading H3K27me3 and that high levels of H3K27me3 are incompatible with high levels of Pol II phosphorylated at serine 5.

In summary, 19 (58%) of the 33 identified interactions are supported by experimental evidence as collected from the literature, showing a direct interaction or involving only one unobserved, additional protein (Supplementary Note S1 and Supplementary Table S2). Our predictions complement the experimental evidence obtained either *in vitro* or by using one or few genes as model system. In addition, as we used ChIP-Seq data the inferred interactions between CMs and HMs provide evidence for the interactions *in vivo* and genome-wide. Finally, we provide testable hypotheses regarding novel interactions, which may be instrumental to define chromatin signaling and its impact on transcription.

### Verification of two predicted interactions links H4K20me1 to Polycomb-mediated repression

Two predicted interactions involve the HM H4K20me1 and CBX2 and EZH2, which are components of PRC1 and 2, respectively. In both cases the interaction is positive suggesting that CBX2 and EZH2 are involved in setting, stabilizing and/or reading H4K20me1.

Given the biochemical properties of CBX2 and EZH2, a role in setting or stabilizing H4K20me1 seems unlikely. However, CBX2 and EZH2 may directly or indirectly bind to H4K20me1. To test the latter possibility, we performed an immunoprecipitation (IP) against H4K20me1 and probed for the presence of CBX2 and EZH2 (Figure 4A). The presence of a positive signal of CBX2 and EZH2 in the H4K20me1 IP and the absence in the control IgG IP suggests that both proteins interact with H4K20me1 Our results are in line with the idea that H4K20me1 is linked to Polycomb-mediated repression by interacting with PRCs 1 and 2.

**Figure 4.**
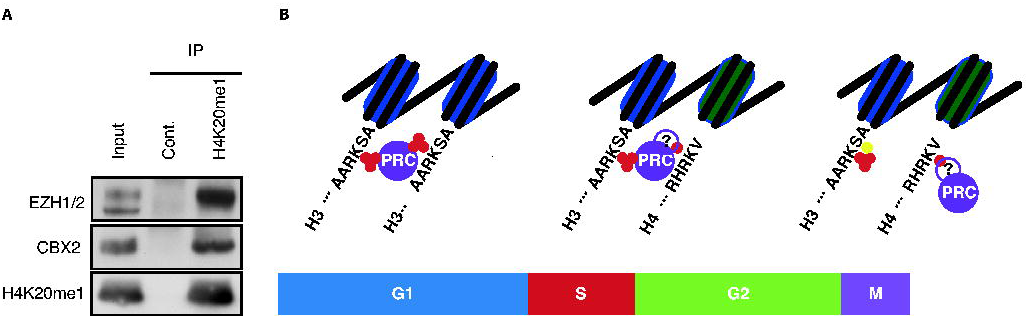
Verification of two predicted interactions links H4K20me1 to Polycomb-mediated repression. (A) H4K20me1 co-immunoprecipitation: K562 cells were immunoprecipitated with a control IgG and an H4K20me1 specific antibody and analysed by immunoblot for the co-precipitation of EZH2, CBX2 and H4K20me1. 10% Input was loaded. (B) Model of the role of H4K20me1 in the maintenance of Polycomb-mediated repression through the cell-cycle. During the G1-phase PRCs bind to H3K27me3 (indicated by three red circles) on two adjacent nucleosomes. During S-phase one of the two nucleosomes is replaced by a new one, which acquires H4K20me1 (indicated by a single red circle). After replication PRCs bind to H3K27me3 on the old nucleosome (in blue) and H4K20me1 on the new (in green), possibly via a yet unknown factor (indicated by the violet circle with the question mark). In M-phase, serine 28 gets phosphorylated (indicated by a yellow circle), which prevents PRCs from binding. PRCs are maintained on chromatin by their interaction with H4K20me1.

## Discussion

Taken together, we propose a novel computational approach to enrich for potential direct interactions linking CMs and HMs within a chromatin-signaling network. We have applied this approach to the most comprehensive set of CMs and HMs in human cells and identified interactions between the CMs and HMs. Furthermore, we have demonstrated that at least two of the predicted but yet unknown interactions can be verified by experimental means. These verified interactions provide an unexplored link between Polycomb-mediated repression and H4K20me1.

Analyzing the pairwise correlation patterns between the levels of CMs and HMs identifies groups of CMs and HMs, which are likely to constitute the building blocks of a chromatin-signaling network. However, unraveling specific interactions between the group members by focusing only on the pairwise correlations is difficult. This difficulty arises from the propagation of correlations along the direct interactions of the network components. For example, H3K27me3 is set by EZH2, which is in a complex with SUZ12 and EED, which itself binds to H3K27me3 (Figure 5A). Thus, the H3K27me3 ChIP-Seq levels correlate with those of EZH2, EED and SUZ12. However, only in the case of EZH2 and EED this is due to a direct interaction with H3K27me3. To remedy such a situation, in our example we need to ask how much more information SUZ12 provides on H3K27me3 given the information provided already by EZH2 and EED. We achieve this by modeling H3K27me3 levels as a weighted linear combination of EZH2, EED and SUZ12 levels. Here, the correlations between EZH2, EED and SUZ12 are taken into account, such that we obtain a weight for SUZ12, which corresponds to the remaining information that SUZ12 has on H3K27me3 after the information of EZH2 and EED on SUZ12 (Figure 5B) and H3K27me3 (Figure 5C) has been subtracted (Figure 5D).

**Figure 5.**
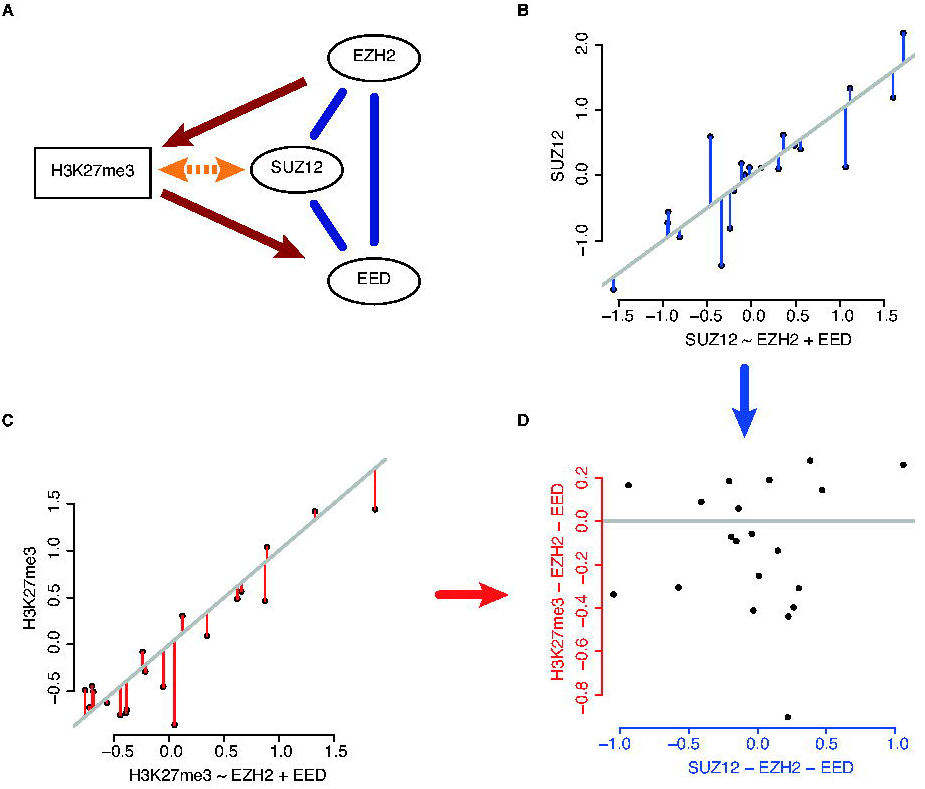
From correlation to direct interactions. (A) Model of the interaction of the PRC2 trimeric complex (EZH2, SUZ12 and EED) with H3K27me3. The blue lines indicate protein-protein interactions. The red arrows indicate direct causal interactions, with EZH2 setting H3K27me3 and EED reading H3K27me3. The orange double-headed arrow indicates a correlation between SUZ12 and H3K27me3 induced by either EZH2 and/or EED. (B – D) Toy example of the decorrelation action of multivariate regression. Modeling of e.g. H3K27me3 levels by a linear combination of EZH2, EED and SUZ12 leads to an estimate of the influence of SUZ12 independent on the influence of EZH2 and EED. This is achieved by modeling SUZ12 (B) and H3K27me3 levels (C) by a linear combination of EZH2 and EED. The predictions of these models are subtracted from the actual SUZ12 and H3K27me3 levels (residuals, depicted by blue (SUZ12) and red (H3K27me3) vertical lines). The residuals of SUZ12 after incorporating the information of EZH2 and EED are used to predict the corresponding residuals of H3K27me3 (D), which in this case fails because there is no information of SUZ12 on H3K27me3 left after considering EZH2 and EED levels.

We use this mathematical framework to explain away indirect interactions and thus to obtain the most direct interactions given the data. This implies that the uncovered interactions may change if additional information is added. For example, we had only data for H3K27me3, EZH2 and SUZ12, but lacked data for EED. Our analysis uncovers an interaction between H3K27me3 and EZH2, which has been shown to set H3K27me3 (18-21). We also identified an interaction between H3K27me3 and SUZ12. The latter interaction is independent of EZH2, but may dependent on the unobserved EED, such that the addition of EED to the dataset will remove the indirect interaction between SUZ12 and H3K27me3.

Within this mathematical framework, we have shown that HM levels are accurately predicted by CM levels and *vice versa* (Figure 2), suggesting a close relationship between CMs and HMs. Given the high predictive power, we are confident to take the weights of the Elastic Net as evidence for an interaction between a HM and a CM. By combining Elastic Net and SPCN we further eliminated indirect interactions moving closer towards a mechanistic understanding of the interactions between HMs and CMs (Figure 3).

These interactions should not be confused with causal interactions. Inference of causality from data requires perturbation experiments as discussed extensively in the literature (29). In our setting such experiments are notoriously difficult to perform, because perturbations of CMs usually either lead to pleiotropic effects, including cell death (30), or are buffered by redundant mechanisms (31, 32). Additionally, manipulation of the histones, i.e. single amino acid substitutions, is not feasible in most organisms except for yeast (33) and Drosophila (34, 35).

Our analysis predicted many interactions between CMs and HMs, of which many are supported by the literature (Supplementary Note S1 and Supplementary Table S2). Others provide novel hypotheses about yet unknown interactions between CMs and HMs, which are amenable to experimental verification. To demonstrate this, we validated two interactions involving the HM H4K20me1 and the CMs EZH2 and CBX2 by co-immunoprecipitation (Figure 4A). These results link H4K20me1 to Polycomb-mediated repression by PRCs 1 and 2, which may form a mechanistic basis for the maintenance of Polycomb-repression through the cell cycle.

The progression of cells through the cell cycle constitutes two challenges for the maintenance of Polycomb-mediated repression: (i) During DNA replication old and newly synthesized nucleosomes are randomly distributed to the daughter strands (36). This leads to an effective dilution of H3K27me3-bearing nucleosomes by half. (ii) During mitosis HMs, chromatin composition and structure change dramatically, rendering the proper transmission of H3K27me3 difficult.

H4K20me1 is tightly regulated during the cell cycle. It starts accumulating during S-phase and attains high levels during mitosis (37). Given this pattern, H4K20me1 may play an important role in maintaining PRCs at their target sites throughout the replicative and mitotic challenges by recruiting PRCs 1 and 2 to regions with old H3K27me3- and new H4K20me1-bearing nucleosomes (Figure 4B).

Taken together, we provide a chromatin-signaling network in K562 cells that links CMs to specific HMs. Our approach aims at high specificity and sacrifices sensitivity leading to high-confidence interactions. We verified two yet unknown interactions, which gives rise to novel biological insights about the interplay between Polycomb-mediated repression and H4K20me1.

## Methods

### ChIP-Seq and gene expression data

The raw HM and CM ChIP-Seq reads were obtained from the SRA Archive (GSE29611 and GSE32509). We merged multiple replicates and mapped uniquely mapping reads to the hg19 genome using Bowtie (38). We counted the number of reads falling into a ±2000bp window centered at the TSSs of all known RefSeq genes (accessed: Oct. 19th, 2012). Only promoter regions with at least one sample having a read count larger than the input control were used. The expression data from Cap Analysis of Gene Expression (CAGE) was obtained from the UCSC genome browser (accessed: Nov. 14th, 2012; K562CellPapAlnRep1/2.bam and H1hescCellPapAlnRep1/2.bam). The CAGE-counts were averaged over the available replicates.

### Read count normalization

We normalized the HM and CM read counts by the following procedure: We estimated the slope of the correlation between the read counts of the sample (S) vs. the read counts of the input control (C) (adding a pseudo-count of 1) by the median (m = median((S + 1)/(C + 1)) of the ratio between the two over all promoters. The read counts were then replaced by the enrichment of the sample over the input normalized by the median (S_norm_ = (S + 1)/(C + 1) * 1/m). This procedure shrinks all the read counts that are highly correlated with input towards zero. The normalized read counts and average CAGE-counts were log-transformed and scaled to have mean zero and standard deviation one.

### Linear regression and regularization using Elastic Nets

We use a combination of computational methods to decipher the chromatin-signaling network as described in the result section. First, we would like to uncover direct interactions between each HM and the CMs taking into account all other CMs at hand. This can be done by predicting each HM from the CMs using linear regression. Linear regression has been applied in various problems for outcome prediction. Here, apart from achieving good prediction accuracy, we are interested in determining the subset of variables (CMs) that is most useful for the prediction. The latter can be obtained with regularized linear regression methods, which, in contrast to simple linear regression models, impose soft constraints on the number of non-zero coefficients. Moreover, it would be desirable that correlated variables, i.e. equally good predictors, have similar weights. This is especially useful for our case, where we might have sets of CMs that interact with an HM only when being in a complex. For this reasons, we used Elastic Nets (14) as implemented in the *glmnet*-package (39) for R (40). The objective function of Elastic Net (as for simple linear regression) is the RSS criterion: 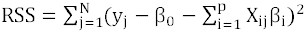 which is the sum of squared errors that should be minimized. In the Elastic Net this objective function is subjected to the constraint: 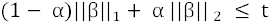 where 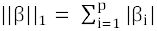 and 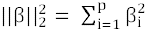 for α E [0,1] and some *t*. The first constraint is based on the L1-norm and forces the coefficients to shrink to 0, thereby favoring sparsity (LASSO-type). The second constraint is based on the L2-norm and favors similar values for the coefficients (Ridge-type), thereby avoiding picking one variable over another when both are redundant. The α -parameter specifies the contribution of each constraint. Throughout the paper we first choose α between 0.01 and 0.99 using 10-fold cross-validation (CV) on each cross-fold. The best α is selected such that the average RSS of the selected α lies within standard deviation of the α having the minimal average RSS. Once α is fixed, the *t*-parameter is then automatically optimized by the *cv.glmnet*-function in a similar fashion.

We estimated the importance of a CM in predicting a specific HM using Elastic Nets and 10-fold CV. Due to the large number of promoters and due to the smoothing operated by the L2-norm, we expect all coefficients to be non-zero, as the prediction accuracy will increase more with one coefficient than the penalty. However, the L1-norm will enhance the contrast between useful and unuseful variables, and will make the selection for the network representation easier. For the graphical representation of the important CMs we select only those CMs that have an average coefficient that deviates from the average of all coefficients by at least one standard deviation (Supplementary Figure S4).

### Partial correlations and the sparse partial correlation network

We combine the Elastic Net approach described above with Sparse Partial Correlation Networks (SPCN) (15), which take into account both HMs and CMs. The SPCN approach is based on the partial correlation coefficient P(X,Y|Z) that gives the correlation coefficient between X and Y after they are controlled for Z. In other words, X and Y are both regressed against the control set Z, and the correlation between their respective residuals r(X) and r(Y) is computed. This allows us to focus on associations that are as direct as possible within the dataset at hand. For a dataset D, the pairwise partial correlations P(X,Y|D\{X,Y}) between every pair X and Y, where all other variables D\{X,Y} are in the control set, can be efficiently computed by inverting and normalizing the covariance matrix of a dataset D.

We build the SPCN on all CMs and HMs(15). In short, we compute the pairwise partial correlation between the ranked ChIP-Seq levels of a CM and an HM conditioned on all other variables (Supplementary Figure S5). Only those edges having a significant, non-zero partial correlation coefficient are retained. Sparseness is introduced in a 10-fold CV scheme which, at the same time, is designed to maintain high accuracy of the resulting (15). For the graphical representation we select only those links from the full SPCN that are between HMs and CMs.

### Cell Lysis and Immunoprecipitation

K562 cells (3x10^6^) were lysed in 350 μl cytoskeletal (CSK) lysis buffer (10mM PIPES, 100 mM NaCl, 300 mM Sucrose, 3mM MgCl_2_, 0.1% NP40) for 10 minutes on ice. The lysate was then centrifuged at 5,000 x g for 5 minutes and the supernatant discarded. The pellet was resuspended in 350 μl of chromatin lysis buffer (300 nM NaCl, 50 mM Hepes pH 7.4, 0.5% Igpal, 2.5 mM MgCl_2_, 5 U Benzonase from Novagene, 1x protease inhibitor cocktail from Roche) for 30 minutes on ice, with periodic mixing. The lysate was centrifuged at 13,000 x g for 10 minutes and the supernatant collected.

2 μg of a mouse IgG control antibody (Diagenode C15200007) or 2 μg of a monoclonal mouse H4K20me1 antibody (Diagenode C15200147) were incubated with 10 μl of magnetic protein G beads (Dynabeads Life Technologies) for 2 hours under rotation at 4°C and then washed several times in the IP buffer. 150 μl of nuclease digested chromatin lysate was diluted with dilution buffer (100 nM NaCl, 50 mM Hepes) to 500 μl and incubated with the antibody coated beads for 4 hours under rotation at 4°C. The beads were then washed 3 x with IP buffer and resuspended in 50 μl of chromatin lysis buffer supplemented with 10 μl of 5x Lammeali buffer. The input and immunoprecipitations were then heated to 99°C for 10 minutes prior to loading on a 4 - 12% gradient gel (Invitrogen). The immunoblot was detected with specific antibodies against H4K420me1 (Abcam ab9057), EZH2 (Epitomics 1940-7) and CBX2 (Abcam ab18968 and Bethyl A302-524A).

## Acknowledgements

This work was supported by EU-FP7 “BLUEPRINT” [282510], Bundesministerium für Bildung und Forschung “Deutsches Epigenom Programm (DEEP)” [01KU1216C] and Deutsche Forschungsgemeinschaft [SFB618].

